# Quantifying Pathological Progression from Single-Cell Data

**DOI:** 10.1101/2024.11.27.625593

**Authors:** Samin Rahman Khan, M. Sohel Rahman, Md. Abul Hassan Samee

## Abstract

The surge in single-cell datasets and reference atlases has enabled the comparison of cell states across conditions, yet a gap persists in quantifying pathological shifts from healthy cell states. To address this gap, we introduce single-cell Pathological Shift Scoring (scPSS) which provides a statistical measure for how much a “query” cell from a diseased sample has been shifted away from a reference group of healthy cells. In scPSS, The distance of a query cell to its *k*-th nearest reference cell is considered as its pathological shift score. Euclidean distances in the top *n* principal component space of the gene expressions are used to measure distances between cells. The p-value of a query pathological shift score belonging to the null distribution of intra-reference cell shift scores provides a statistical significance measure of the query cell being in the reference cell group. This makes our method both simple and statistically rigorous. Comparative evaluations against a state-of-the-art contrastive variational inference model, modified for shift scores, demonstrate our method’s accuracy and efficiency. Additionally, we have also shown that the aggregation of cell-level pathological scores from scPSS can be used to predict health conditions at the individual level.

## Introduction

The ability to measure cellular state transitions between healthy and diseased conditions is fundamental to understanding disease mechanisms and progression. The increasing availability of single-cell datasets and large-scale reference atlases (Lindeboom et al., 2021; Regev et al., 2017) enables comparing cell states across different conditions (Nomura, 2021). However, existing tools lack the capability to identify cell populations that have shifted statistically significantly from a reference state - a critical aspect for accurate disease characterization. To address this need, we introduce single-cell Pathological Shift Scoring (scPSS), a computational method that quantifies the statistical significance of cellular state deviations from a reference. Importantly, scPSS does not require annotated disease datasets, making it uniquely applicable to rare and emerging diseases.

Current methods based on dimensionality reduction (Kobak & Berens, 2019; Xiang et al., 2021) and machine learning (Lotfollahi et al., 2021; Michielsen et al., 2023) can effectively classify cells into known states but cannot quantify the degree of shift from a reference state. Linear models using principal component (PC) embeddings of gene expression are popular for single-cell analysis, with distances in PC space being used to cluster cells into groups (Fa et al., 2021) and measure differences between these groups (Nicol et al., 2023). Furthermore, contrastive methods (Abid et al., 2018; Gorla et al., 2023) and contrastive learning approaches (Weinberger et al., 2023) have been developed to identify specific features that are more informative in distinguishing between cell groups through targeted dimensionality reduction. While these approaches have improved disease-specific feature identification, they don’t provide a quantitative assessment of state alterations and lack the ability to measure the degree of disease-associated changes. Recent machine learning methods have enabled the identification (Goeva et al., 2024) and scoring (Litinetskaya et al., 2024) of disease-relevant cellular states, but their dependence on labeled training data from both healthy and diseased individuals limits their application to well-characterized conditions.

Our approach, scPSS, uses gene expression profiles from normal cells to establish a reference state distribution, using k-nearest neighbor distances in principal component embedding space. For any query cell, it calculates a “pathological shift score” that measures its deviation from this healthy state distribution. This score enables both the ranking of cells by their degree of state deviation and the identification of disease progression. The key differentiator for scPSS is its “zero-shot” capability - quantifying pathological shifts using only healthy reference distributions, without requiring training on condition-labeled data. By employing a simple yet robust statistical framework, scPSS provides an interpretable measure of cellular state deviation while adhering to the principle of Occam’s Razor.

In comparative analyses across multiple datasets containing healthy and diseased cells, scPSS matched or exceeded the performance of the state-of-the-art Contrastive VI method (Weinberger et al., 2023) (modified to provide pathological shift scores), showing substantial improvements in AUPR of up to 32% in most datasets **(Table 1)**. We also used the scPSS to identify healthy and diseased individuals from the proportion of diseased cells compared with reference datasets. Using the HCLA dataset, scPSS successfully distinguished healthy from IPF conditions with 86% accuracy using only healthy reference cells, and 91% accuracy when both healthy and diseased reference cells were available.

**Table 1.** Comparison of AUC and AUPR measures between scPSS (our method) and the adapted ContrastiveVI model for predicting healthy and damaged cells from using pre-MI (Myocardial Infarction) dataset as reference and cells collected at different time points after MI as query dataset. Results show mean ± 95% confidence interval from 25 independent runs for scPSS and 5 runs for ContrastiveVI.

| Query dataset | AUC |  | AUPR |  |
| --- | --- | --- | --- | --- |
|  | Contrastive VI (adapted) | scPSS (Ours) | Contrastive VI (adapted) | scPSS (Ours) |
| 1HR | 0.8180 ± 0.1492 | **0.9107 ± 0.0022** | 0.7271 ± 0.2042 | **0.8452 ± 0.0037** |
| 4HR | 0.7487 ± 0.3376 | **0.9478 ± 0.0011** | 0.8126 ± 0.2534 | **0.9649 ± 0.0007** |
| snd1 | 0.7609 ± 0.1852 | **0.9507 ± 0.0004** | 0.8098 ± 0.1473 | **0.9590 ± 0.0003** |
| snd3 | 0.7614 ± 0.1320 | **0.9282 ± 0.0007** | 0.6784 ± 0.1447 | **0.8952 ± 0.0009** |
| snd7 | 0.6893 ± 0.0104 | **0.8476 ± 0.0017** | 0.6243 ± 0.0087 | **0.8185 ± 0.0028** |
| all post-MI | 0.7686 ± 0.0578 | **0.9009 ± 0.0006** | 0.7626 ± 0.0581 | **0.9023 ± 0.0005** |

## Results

### Overview of scPSS

We find the pathological shift score of a query cell using the distance to its k-th nearest reference cell. The Euclidean distances between cells are measured in a lower dimensional embedding of the gene expression values in order to better capture the latent biological representations. We have used principal component analysis to get the latent representations. As a preprocessing step, the principal component projection values were adjusted using Harmony (Korsunsky et al., 2019) to account for the batch-specific differences between the different datasets. For the statistical significance of the pathological shift scores we first prepare a null distribution of distances of reference cells from their k-th nearest neighbour. Then, the p-value of the distance of the query cells from their k-th nearest neighbor, i.e., their pathological shift score belonging to the null distribution is determined. **Fig. 1** shows the scPSS pipeline. Further details can be found in the Methods Section.

**Figure 1.**
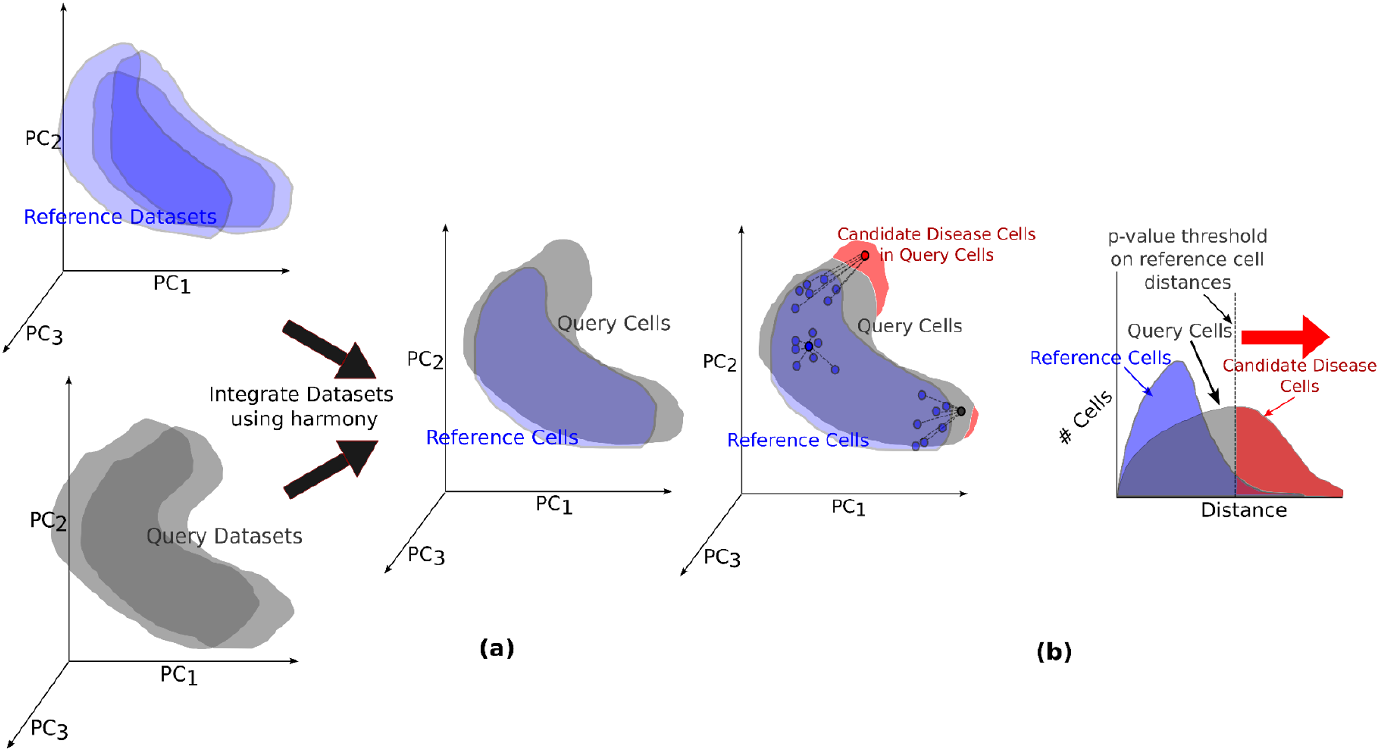
Overview of scPSS: **(a)** At first, principal component analysis is done on individual query and reference datasets. These individual datasets are integrated using the harmony method. This removes batch-specific effects on these datasets by adjusting the principal component (PC) values. **(b)** The Euclidean distance of a cell to the k-th nearest reference cell in PC space is used as its pathological shift score. The distances of the reference cells with their neighboring reference cells are considered as the null distribution. We can determine the p-value of each query cell distance belonging to the reference null distribution to get a significance measure for pathological progression.

### scPSS Identifies Damaged Cells and Damage Progression in Infarcted Heart Tissue of Mouse

We validated scPSS using single-cell transcriptomic data from mouse hearts before and after myocardial infarction (MI) (Calcagno et al., 2022). The dataset contains labeled cardiomyocytes (CMs) from three distinct regions: the remote zone (RZ, far from infarction), border zone 1 (BZ1), and border zone 2 (BZ2, both adjacent to infarction) **(Fig. 2a)**. Pre-MI samples predominantly contain RZ cells, while post-MI samples include a mixture of RZ, BZ1, and BZ2 cells, providing an ideal benchmark for testing how well different methods predict outlier disease cells using healthy cells as reference **(Fig. 2b)**.

**Figure 2.**
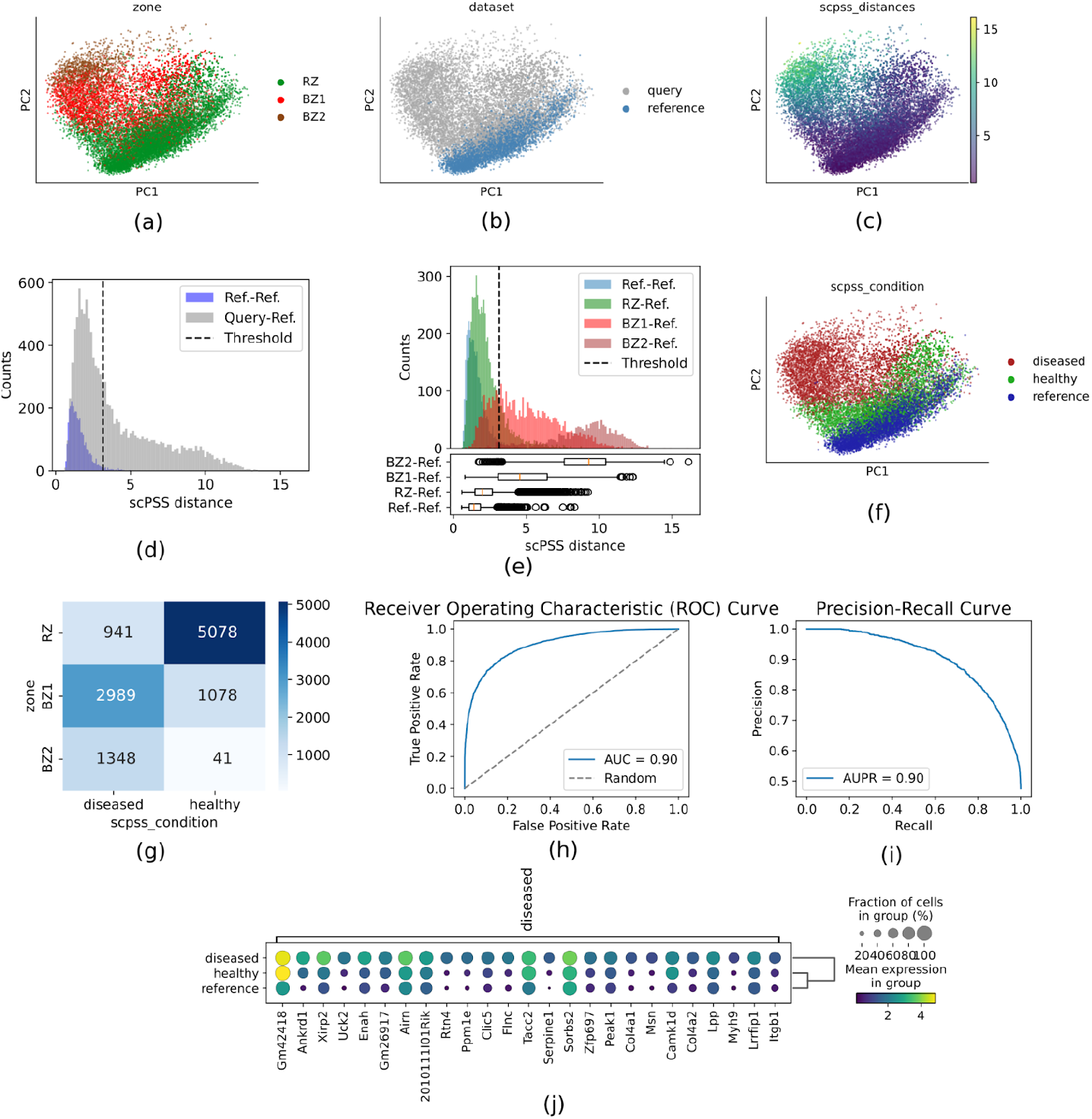
scPSS for damage progression in Infarcted Heart Tissue. **a-c**. The PC embeddings of all the cells from both healthy (before infarction) and query (after infarction), colored according to regions of nearness to infarcted regions (a), according to whether they belong to healthy or query dataset (b) and the shift distance assigned by scPSS (c). **d**. Distance distribution of reference cells with reference cells and query cells with reference cells. **e**. Distance distribution of reference cells with reference cells and remote zone (RZ), border zone 1 (BZ1), and border zone 2 (BZ2) query cells with reference cells in histograms and boxplots. **f**. The PC embeddings of all the cells from both healthy (before infarction) and query (after infarction), colored according to disease condition labels provided by scPSS **g**. The confusion matrix of the true labels and predicted labels given by scPSS framework on the query cells. **h-i**. The Reciever Operating Characteristic (ROC) (h) and Precision-Recall (i) curves for the pathological scores provided by our method on the query dataset. **j**. Top 25 differentially expressed genes across cells predicted diseased and healthy from query dataset and reference cells.

For our benchmark test, we used pre-MI (day 0) samples as the reference dataset and post-MI samples (collected at 1hr, 4hr, 1 day, 3 days, 7 days, and all post-MI time points) as query datasets, with RZ cells representing the healthy state and BZ1 and BZ2 cells representing the disease state. We compared scPSS against the modified ContrastiveVI model using both Area Under the Receiver Operating Characteristic curve (AUC) and Area Under the Precision-Recall curve (AUPR) metrics **(Table 1) (Fig 2h-i)**. scPSS outperformed ContrastiveVI in all time points, demonstrating its superior ability to rank cells by their deviation from the healthy reference state.

An optimal threshold is determined that best separates the reference and query distributions to classify query cells into healthy and diseased **(Methods) (Fig. 2d, f)**. For the case where pre-MI cells are used as reference and post-MI cells at all time points as query, scPSS was able to classify RZ cells as healthy and BZ1 and BZ2 cells as disease at an accuracy of ∼80%. We also found that the distances of BZ2 from the reference were more than the distances of BZ1 **(Fig 2e)**. Furthermore, this classification of query cells allowed us to identify disease-associated genes through differential expression analysis between the predicted diseased and reference cell populations **(Fig. 2j)**.

### scPSS Classifies the Condition of Individuals from Pathological Cell Proportions

We evaluated scPSS’s ability to classify disease states at the individual level using a subset of the Human Lung Cell Atlas (HLCA) (Sikkema et al., 2023) that has been used in the study by Litinetskaya et al. (Litinetskaya et al., 2024), containing single-cell data from 59 healthy individuals and 52 patients with idiopathic pulmonary fibrosis (IPF). Using the pathological progression scores of each individual cell it is also possible to provide the pathological labels of each individual organism/specimen. scPSS provides a pathological distance for the query cells and a threshold above which a query cell can be considered as pathological **(Fig. 3c)**. The individual is labeled as healthy or diseased depending on the proportion of pathological cells from a cell type in an individual. Individuals in the query dataset with significantly higher disease proportions than reference healthy individuals are considered to be diseased **(Fig. 3d) (Methods)**.

**Figure 3.**
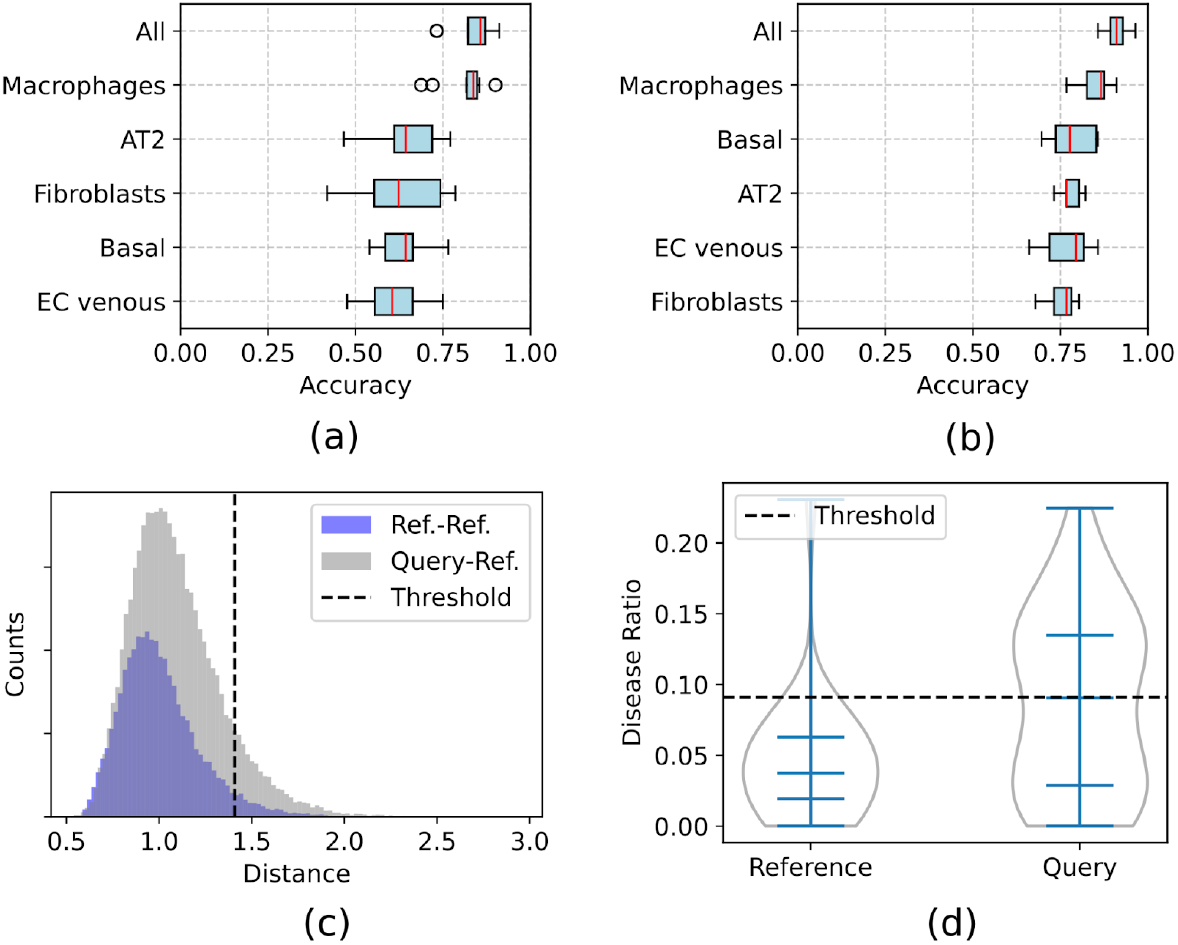
scPSS predicting healthy vs IPF. **a**. With only healthy cells present in the reference dataset, the accuracy of scPSS pipelines to predict the health vs IPF condition of individuals using the proportion of pathological cells of each cell group and all cells together. **b**. With cells from both healthy and diseased individuals in the reference dataset, we see an increase in predictive accuracy for the same experiment. (*Note:* for calculating accuracy for a certain cell type, we considered only the individuals with cells of that type.) **c**. Distance distribution of reference cells with reference cells and query cells with reference cells for the Macrophage cell type in one example fold. **d**. Violin plots outlier ratios or disease ratios of reference and query individuals for the Macrophage cell type in one example fold.

In our initial experiment, we used healthy reference data only, randomly selecting 50% of healthy individuals as reference and the remaining healthy individuals plus 50% of IPF patients as query samples. This problem setup reflects the scenario when we have cell samples from healthy individuals as reference and need to classify individuals as healthy and diseased. To ensure robust evaluation, we repeated this procedure 10 times with different random choices of individuals into reference and query sets. Analysis of individual cell types revealed that Macrophages alone achieved a median classification accuracy of ∼84%. Other cell types, including EC Venous, Fibroblasts, Basal, and AT2 clusters, also proved to be strong indicators of disease state. Considering the disease portions of these five cell types together **(Methods)** yielded a median accuracy of ∼86% **(Fig 3a)**.

If we have both healthy and diseased individuals in our reference dataset, we can better predict healthy and diseased individuals in the query dataset **(Methods)**. To show this, we repeated the same experiment. But this time, we randomly picked 50% of the healthy individuals and 50% of the diseased individuals into the reference datasets and the rest into the query dataset. We have found the accuracy in this case using indicative cell types to be ∼91% **(Fig 3b)**.

### scPSS Identifies Pathological Progression in BMMC of Leukemia Patients

We applied scPSS to analyze bone marrow mononuclear cells (BMMC) from leukemia patients before and after bone marrow transplantation (Zheng et al., 2017). From this dataset, we have used cells from healthy patients as our reference and cells from patients as our query datasets to check the pathological progression of cells in pre-transplant and post-transplant conditions. Pathological shift scores were calculated using the distance to the 270th nearest reference cell (see **Methods** for parameter selection). scPSS showed that cells from pre-transplant patients (67% with a p-value of < 0.05) were more pathological than those from post-transplant cells (37% with a p-value of < 0.05). This reduction in pathological cells following transplantation aligns with expected clinical recovery patterns and demonstrates scPSS’s utility in monitoring treatment response.

## Discussion

scPSS provides a fast but accurate statistical method for finding pathological shifts of cells from a reference cell by using distance measures in the principal component space of gene expression values.

Distinguishing biological signals from technical artifacts remains a key problem in single-cell analysis, and scPSS is no exception. Batch effects, present in most datasets, can obscure changes that are disease-causing. scPSS employs harmony for batch effect removal. However, this process may potentially remove some disease-related variation along with the technical noise, particularly when the proportion of diseased cells is unknown. Determining the cut-off point for classifying the query cells as diseased based on the pathological shift score is still a choice left up to the user of scPSS. However, scPSS provides good heuristic approaches in choosing such parameters. There may be a complex non-linear mapping between genotypical and phenotypical space, which cannot be always captured by the linear PC embeddings.

Despite the limitations, scPSS provides a good benchmark for finding pathological progression. This has been made apparent by comparing scPSS with the best existing models on standard query and reference datasets. Our results demonstrated scPSS’s effectiveness across multiple disease contexts. The method’s ability to capture meaningful biological transitions was validated by its accurate ranking of disease progression in infarcted heart tissue (BZ1 and BZ2) and its detection of healing patterns in bone marrow transplant patients. Furthermore, scPSS successfully translated these cellular-level assessments to individual-level disease classification, as demonstrated in the IPF study where analysis of pathological cell proportions enabled accurate patient diagnosis. These findings establish scPSS as a robust tool for quantifying disease progression at both cellular and organism levels.

## Data and Code Availability

Datasets (Calcagno et al., 2022), (Sikkema et al., 2023) and (Zheng et al., 2017) used in this study are publicly available at https://www.ncbi.nlm.nih.gov/geo/query/acc.cgi?acc=GSE214611, https://drive.google.com/uc?export=download&id=1wWGwbPeap-IqWNVlwVVUWVrUAMrf45ye and https://www.10xgenomics.com/datasets respectively. The code for scPSS is available at https://github.com/SaminRK/scPSS and the code to download all datasets and reproduce all the results for this study is shared at https://github.com/SaminRK/scPSS-reproducibility.

## Methods

Suppose, *R* be the set of reference cells and *Q* be the set of query cells. The gene expression values of *g* genes of the reference and query cells can be represented as *X* ∈ ℝ^*m*×*g*^, where *m* = |*R*| + |*Q*|. The goal is to find the pathological shift distances, Δ ∈ ℝ^|*Q*|^, for the query cells with respect to the reference cells and the statistical significances, *P* ∈ [0, 1]^|*Q*|^, of these shift distances. In addition, we also want to find the pathological labels, *Pr* ∈ {0, 1}^|*Q*|^, of the query cells, where diseased cells are labeled as 1 and healthy cells are labeled as 0. We consider that the reference cells *R* and query cells *Q* are of the same cell type cluster. If there are multiple cell types, we only keep cells of a single type which is of interest in *R* and *Q*.

### Integrating Reference and Query Single-Cell Datasets

The principal components of the gene expression values of the cells in the combined dataset *R* ∪ *Q* are found. Let *PC*: ℝ^*g*^ → ℝ^*N*^ be the PCA transformation that maps the gene expression values of the cells to their *N*-dimensional PC embeddings, where *N* < *g*. The principal component projection scores are adjusted using the Harmony (Korsunsky et al., 2019) method to remove batch-specific effects from the datasets. Let *P*: ℝ^*N*^ → ℝ^*N*^ be the Harmony transformation. Harmony more reliably preserves biological signals as compared to other batch correction methods (Antonsson & Melsted, 2024). Harmony does soft clustering on PC space to group cells according to their cell types or states. Correction factors for each dataset for each cell type or state are found using the cell type or state-specific centroids. Then the PC values of the individual cells are adjusted by their correction measures. The clustering and correction stages are done iteratively until convergence.

### Calculating Pathological Shift Scores

We measure distances between cells in the adjusted principal component space. We use the top *n* principal components to measure these distances. The key genes driving differences in cell state will likely be included in the top principal components, as they account for a major portion of the variance between inlier and outlier cells. Let *P* ∈ ℝ^*m*×*n*^ represent the top adjusted PC embeddings of the cells, where *n* is the total number of top principal components used. The distance between two cells *i* and *j* is defined as,

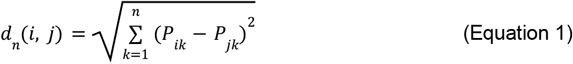

We then use k-nearest neighbor outlier detection(Ramaswamy et al., 2000) to calculate the pathological shift scores. Specifically, for each cell *q*, we find the distance to its k-th nearest neighboring reference cell *NN*_*k*_(*q*). These distances are considered the pathological shift scores for each cell. The formula for the shift distance can be defined as,

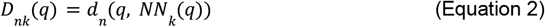

The value of *n*, the number of principal components used, and the choice of *k* for the k-nearest neighbor approach are chosen based on the characteristics of the dataset by searching for the best parameters (discussed below).

### Statistical Significance of Pathological Shift Scores

To assess the statistical significance of pathological shift scores, we first construct a null distribution using the k-th nearest neighbor distances among reference cells. To ensure a continuous p-value measure for query distances, we fit a continuous probability distribution function to these reference distances. We compared different distribution functions (gamma, log-normal, Weibull, and exponential) using goodness-of-fit as assessed using Kolmogorov-Smirnov tests (Massey, 1951) as shown in **Supplementary Figures 1 and 2** and found log-normal and gamma distributions to be the best. The formula for probability distribution function of the gamma distribution is

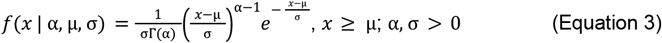

where α is the shape parameter, µ is the location parameter, σ is the scale parameter and is the gamma function defined by the formula

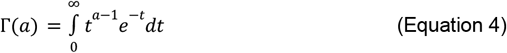

The formula for probability distribution function of the log-normal distribution is

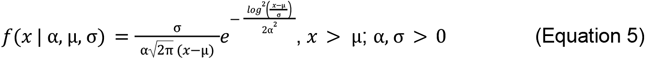

where α is the shape parameter, µ is the location parameter and σ is the scale parameter.

Parameters α, µ, σ of the distribution function *f* are estimated using maximum likelihood estimation using Scipy’s implementation (Virtanen et al., 2020). For each query cell, we compute its pathological shift score and calculate its p-value against this fitted null distribution. If *f*(*x* | α, µ, σ) is the probability density function and 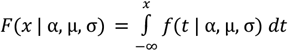 is the corresponding cumulative density function, then the statistical significance (p-value) of a shift distance Δ can be measured using,

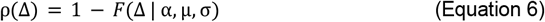

A low p-value would suggest that the corresponding query cell has shifted away significantly from the reference cells and is an outlier. A threshold p-value *p* (such as <0.05) can be picked below which all query cells are considered diseased.

### Parameter selection for scPSS framework

For labeling cells as diseased in a query dataset, the following parameters are to be set for the scPSS framework: (1) *n*, the number of principal components used for distance calculations, (2) *k*, specifying which nearest neighbor distance to use for the pathological shift score, and (3) *p*, the significance threshold below which cells are classified as pathological. We want to pick parameters that differentiate the most number of query cells from the reference cells. So, we choose parameters that provide greater outlier ratios (proportion of query cells labeled as outliers) for a certain threshold.

For selected parameters n, k, and p, the outlier ratio can be defined as below,

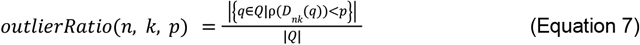

For parameter optimization, we first determine the optimal k value by evaluating outlier ratios (proportion of cells classified as pathological) across multiple significance thresholds (p = 0.01, 0.05, and 0.1) in a given n-dimensional PC space. The k value yielding the highest mean outlier ratio across these thresholds is selected.

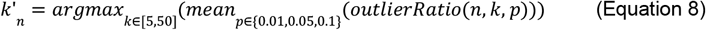

Using this optimal *k*’_*n*_, we then generate a curve of outlier ratios for p-values ranging from 0.01 to 0.15. The optimal p-value threshold is determined using the Kneedle algorithm (Satopaa et al., 2011) to identify the point of diminishing returns in this curve.

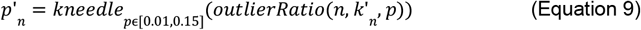

This process is repeated across different values of n (typically 2-20 principal components) to identify the dimensionality that achieves the maximum outlier ratio.

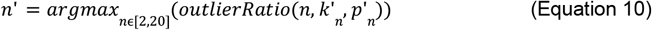

If the reference dataset contains both healthy and diseased cells, the level of integration becomes an additional controllable parameter. Insufficient integration may leave batch effects unresolved, potentially increasing the number of detected outliers. On the other hand, excessive integration can obscure disease-specific effects, compromising outlier detection. The integration level can be controlled by adjusting the maximum number of iterations in the Harmony integration process. Without the presence of diseased individuals in the reference, we do not have any labeled healthy and diseased cells through which we can pick the optimal integration parameters. So, we pick the default one provided by Harmony, which is 10. However, when diseased cells are present in the reference dataset, we can optimize the integration by running scPSS on the reference cells alone. We continue Harmony integration until we observe a drop in the outlier ratio in diseased individuals from the reference dataset compared to healthy individuals, as shown in **Supplementary Figure 3**. It is to be noted that in this case we are not using the query cells, but only the reference, to determine the parameter of maximum number of iterations for Harmony.

### Determining the Pathological Condition of Query Cells

When disease-labeled cells are absent in the reference set *R*, we label a query cell *q* as pathological as *Pr*(*q*) = 1 if ρ(*D* (*q*)) < *p*’ else 0.

When disease-labeled cells are present we find the optimal parameters, query cells are labeled using k-nearest neighbors classification (Cover & Hart, 1967). We first find the optimal parameters *n*’, *k*’, and *p*’ considering reference *R*’ = {*r* ∈ *R* | *r* is healthy} and query *Q*’ = {*r* ∈ *R* | *r* is diseased}. We classify the reference cells from diseased individuals as *Pr*(*r*) = 1 if ρ(*D n*’*k*’ (*r*)) < *p*’ else 0. Using the labeled reference and diseased cells from the reference, the original query cells *q* ∈ *Q* are classified by k-nearest neighbors classification (k=3).

### Aggregating the Pathological Scores of Different Cell Types to Predict Pathological Condition of Individual

To classify the disease condition of an individual, we first analyze each cell type separately using scPSS to calculate the proportion of diseased cells. Let *Q*_*Ii*_ be the query cells of cell type *i* of an individual *I*. The proportion of diseased cells of the cell type *i* of the individual *I* can be determined using

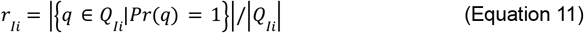

We then combine these proportions across different cell types using one of several aggregation methods to make a final disease classification. The following strategies have been applied for aggregating the disease proportion of different cell types to provide the final label for the individual:

- **Majority Voting:** For each cell type *i* that is considered for disease labeling, we first find the upper limit threshold of disease proportion θ_*i*_ based on disease proportions observed in healthy individuals. We first identify potential outliers using the standard box plot method (values above *Q*_3_ + 1. 5 × *IQR*) (Tukey, 1977). The upper limit is then set as the highest observed value that isn’t considered an outlier. For a cell type *i*, the disease proportion *r*_*i*_ can be considered disease indicating if *r*_*i*_ > θ_*i*_, otherwise it can be regarded as normal. If there are more cell types indicating diseased condition compared to normal condition, the individual is labeled as diseased.
- **Average Disease Proportion:** This method compares the disease proportions of all considered cell types and compares them to reference thresholds. For each cell type *i*, we calculate the difference between the observed disease proportion, *r*_*i*_, and its cell-type-specific upper limit threshold, θ_*i*_. Mathematically, the aggregate score Ψ is defined as: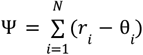. An individual is classified as diseased if Ψ > τ, where τ is the classification threshold.
- **Adjusted Average Disease Proportion:** In this method, the disease proportions of cells of each cell type are normalized as follows: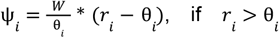, else 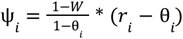. We label the individual as diseased if the aggregate score 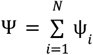 is greater than τ, where τ is the classification threshold.

### Using ContrastiveVI to Find Pathological Scores

ContrastiveVI (Weinberger et al., 2023) is a deep learning model that identifies salient latent features present in a target dataset but absent in a reference dataset. The query cell can then be clustered into groups based on these salient features. We have repurposed this model to find the amount of shift of query cells from the reference cells in order to see how our approach fairs against it. Since disease-specific features are only available in the query dataset, the disease-specific features can be captured by the ContrastiveVI model. Using these disease-specific features, the cells of the query cells can be separated into healthy and disease clusters.

The model is trained by using the reference dataset as the background and the query dataset as the target. The salient latent feature values of query cells were found. We then applied k-means clustering (k=2) to partition the query cells in this latent space. The cluster which was closer to the healthy cells in the salient latent space was considered as the healthy cluster and the other was considered as the diseased cluster. The distances of the cells to the healthy and disease cluster centers were used as a measure of shift, according to the following formula, shift score of cell, *S* = *d*(*C, C*_*d*_) − *d*(*C, C*_*h*_). Here *d* is the Euclidean distance measured in the salient latent space, *C* which is the latent representation of the query cell, *C*_*d*_ the disease cluster centroid, and *C*_*h*_ the healthy cluster centroid.

### scPSS on the BMMC of Leukemia Patients Dataset

We have applied scPSS on BMMC single-cell data (Zheng et al., 2017) to measure the pathological progression of single cells from leukemia patients taken before and after the transplant of bone marrow with reference to those from healthy donors. We have taken only the top 2 principal components. In this space, the distance of the 270-th nearest reference cell was used as the score. The value of k=270 for k-th nearest distance was chosen because of the sharp rise in the outlier ratio at k=270 for different p-value cutoff thresholds. With k fixed at 270, a p-value cutoff of 0.05 was chosen by observing the sharp increase of the outlier ratio for p-value cut-offs up to around 0.05 and its diminishing increase with the further increase of p-value cutoffs (shown in **Supplementary Figure 4**).

## Author contributions

Conceptualization, M.A.H.S. and S.R.K.; Methodology, S.R.K, M.A.H.S. and M.S.R.; Software, S.R.K.; Formal Analysis, S.R.K.; Investigation, S.R.K.; Writing - Original Draft, S.R.K; Writing - Review & Editing, M.A.H.S., S.R.K and M.S.R.; Supervision, M.A.H.S and M.S.R.

## Supplemental information

Document S1. Supplementary Figures 1–4

