## Supplementary Document S1 for "Quantifying Pathological Progression from Single-Cell Data"

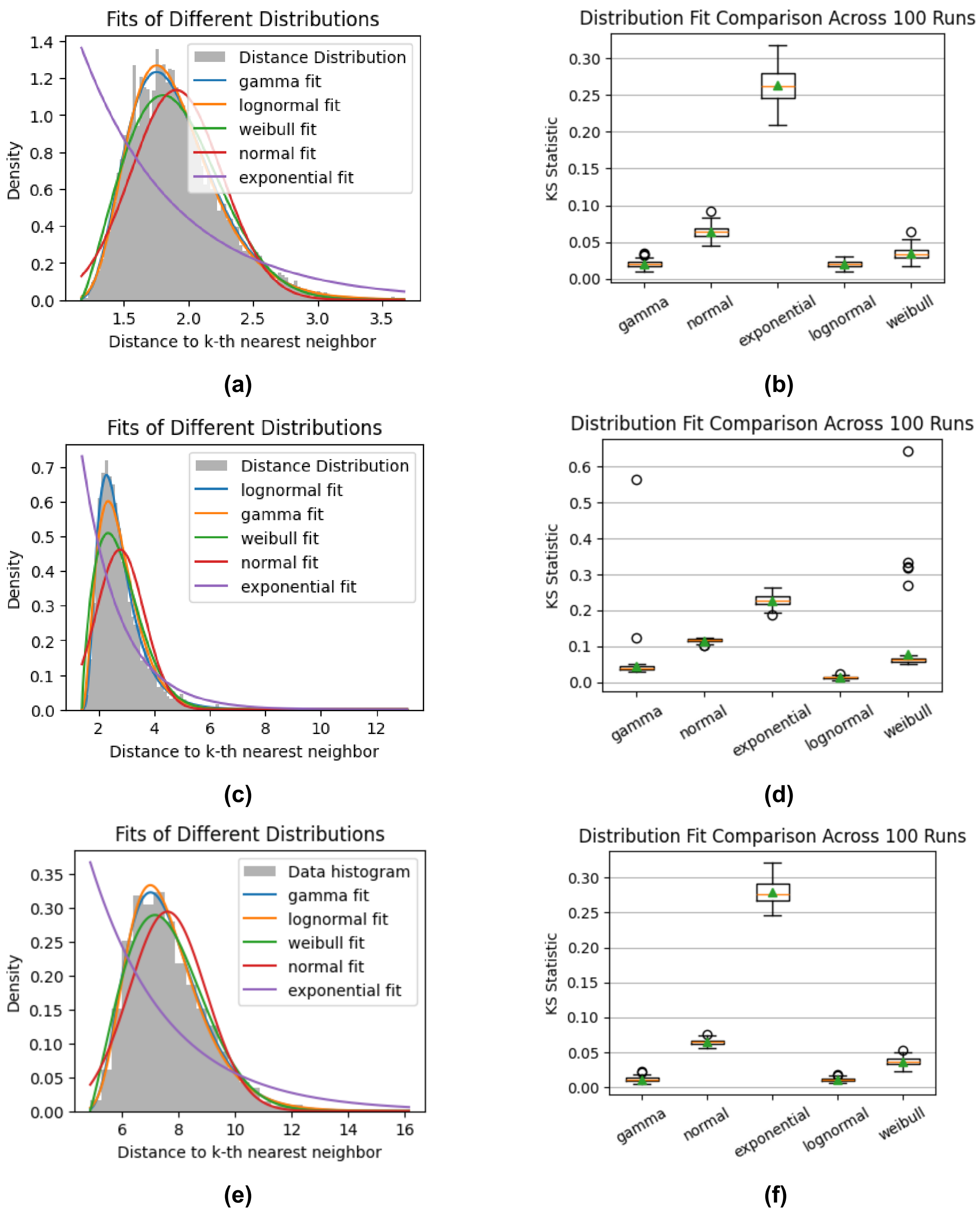


**Supplementary Figure 1:** Goodness-of-fit analysis of various probability distribution functions to a distribution of distances to the 5th nearest neighbor of 5,000 random points in 10-dimensional space **(a)** Distance distribution and fitted distribution functions when points are drawn from $N(0, 1)$. **(b)** Boxplot of KS statistic when this is repeated 100 times. (Median values are gamma: 0.020, lognormal: 0.020, Weibull: 0.034, normal: 0.065, and exponential: 0.262.) **(c)** Distance distribution and fitted distribution functions when points are drawn from $\mathrm{lognormal}(1,\ 0.5)$. **(d)** Boxplot of KS statistic when this is repeated 100 times. (Median values are lognormal: 0.012, gamma: 0.039, Weibull: 0.062, normal: 0.116, and exponential: 0.227.) **(c)** Distance distribution and fitted distribution functions when points are drawn from $N(0, 4)\ +\ N(2, 4)$. **(d)** Boxplot of KS statistic when this is repeated 100 times. (Median values are gamma: 0.011, lognormal: 0.011, Weibull: 0.037, normal: 0.065, and exponential: 0.276.)

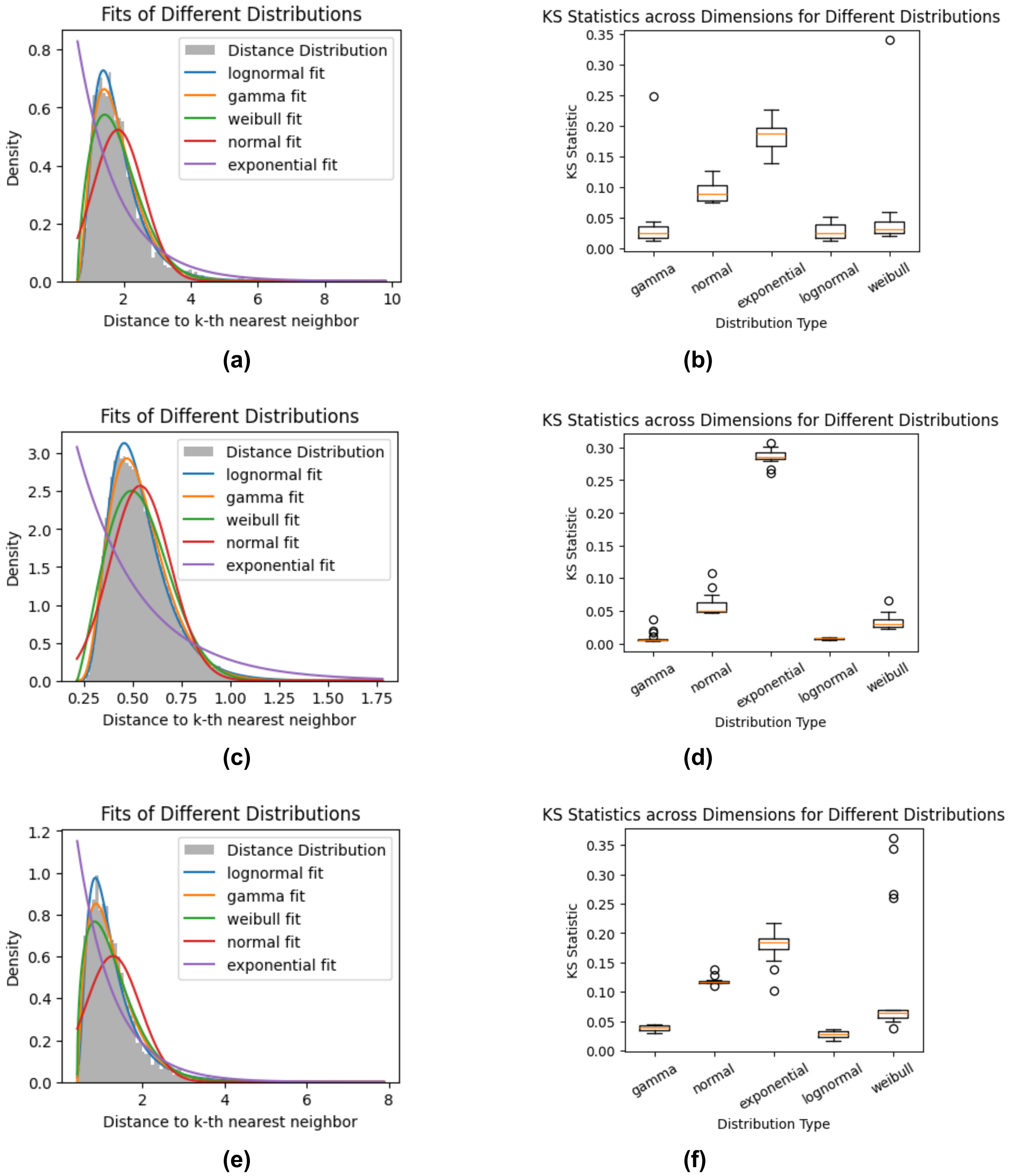


**Supplementary Figure 2.** Goodness-of-fit analysis of various probability distribution functions to a distribution of distances to the 5th nearest neighbor of principal component (PC) embeddings of single-cell transcriptomic datasets **(a)** Distance distribution and fitted distribution functions when top 6 PC embeddings of remote zone Cardiomyocyte cells from Calcagno et al (Calcagno et al., 2022) are used. **(b)** Boxplot of KS statistic when this is repeated with top 4 to 20 top PC embeddings (Median values are gamma: 0.038, lognormal: 0.028, Weibull: 0.051, normal: 0.092, and exponential: 0.182.) **(c)** Distance distribution and fitted distribution functions when top 6 PC embeddings of healthy Macrophage cells from Sikkema et al. (Sikkema et al., 2023) are used. **(d)** Boxplot of KS statistic when this is repeated with top 4 to 20 top PC embeddings (Median values are gamma: 0.009, lognormal: 0.008, Weibull: 0.033, normal: 0.058, and exponential: 0.287.) **(c)** Distance distribution and fitted distribution functions when top 6 PC embeddings of healthy cells from Zhang et al. (Zheng et al., 2017)  are used. **(d)** Boxplot of KS statistic when this is repeated with top 4 to 20 top PC embeddings (Median values are gamma: 0.039, lognormal: 0.027, Weibull: 0.117, normal: 0.119, and exponential: 0.177.)

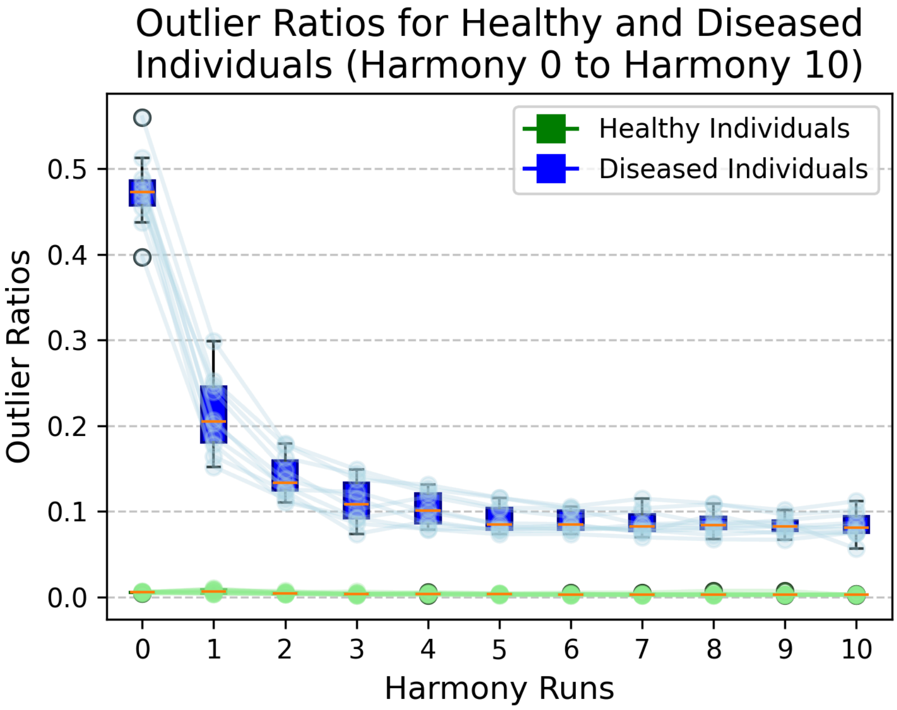


**Supplementary Figure 3.** Increasing the number of iterations of Harmony integration reduces the outlier ratio in diseased individuals. Insufficient integration may leave batch effects unresolved, potentially increasing the number of detected outliers. On the other hand, excessive integration can remove disease-specific effects, compromising outlier detection. When condition-labeled single-cell transcriptomics data from both healthy and diseased individuals were available in the reference, the level of integration can be set as a parameter. This was tested in the HLCA dataset (Sikkema et al., 2023) in the scenario. For all the 10 test runs, on no integration, the outlier ratio of diseased individuals is ~0.5 and after the 2nd iteration of Harmony, it plateaus to ~0.1. On the 1st iteration, the outlier ratio is ~0.2 which is a good balance between no integration and full integration. So, for all the 10 test cases where diseased and healthy labeled data were available, Harmony integration was run for a maximum of 1 time.

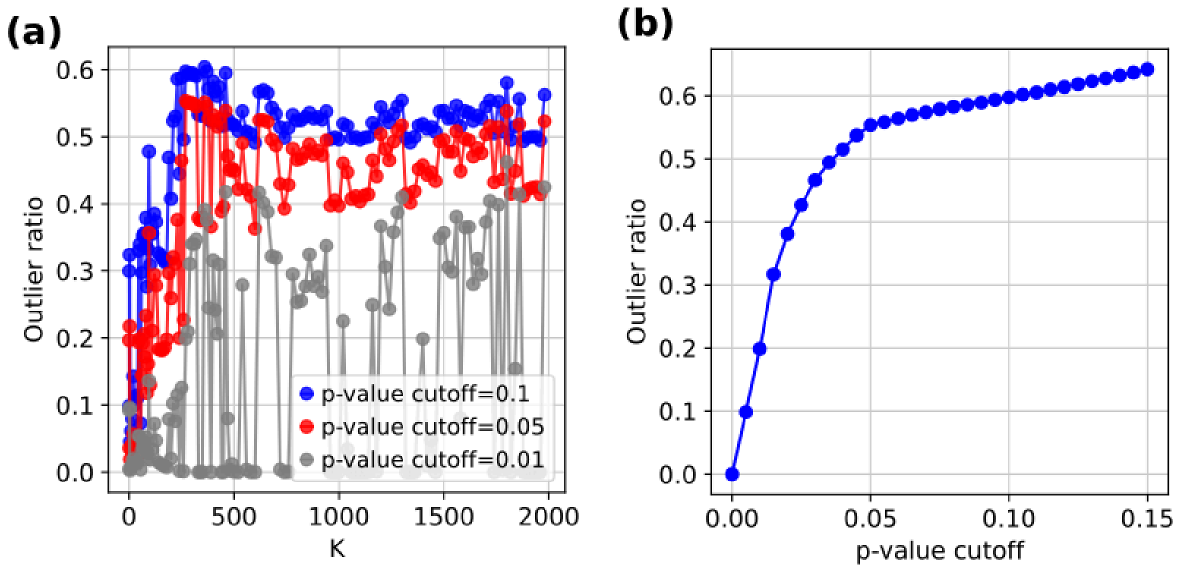


**Supplementary Figure 4. scPSS parameter selection on the BMMC of Leukaemia Patients Dataset. (a)** The curve shows the outlier ratios at different values of parameter k for different p-value thresholds. **(b)** The curve shows outlier ratios at different p-value thresholds for a fixed parameter k. (Distance to k-th nearest neighbor is used as the shift score.)
